# Cutting Through the Artifacts: Dissecting gRNA Impurities with FUSS-seq

**DOI:** 10.1101/2025.09.13.676056

**Authors:** Antonino Montalbano, Huan Qiu, Lakshmi Karthik, Sawyer Letourneau, Jianxin Hu, Tirtha Chakraborty, Huanying Gary Ge, John R Lydeard, Ruijia Wang, Eric Gunnar Anderson

## Abstract

CRISPR-based therapeutics rely on guide RNAs (gRNAs) and the Cas9 endonuclease for precise gene editing. Ensuring gRNA purity and base-level sequence integrity is essential for clinical translation. While industry-standard practice relies on liquid chromatography-high-resolution mass spectrometry to assess oligonucleotide identity and purity, more recent FDA guidance recommends complementary base-by-base sequence analysis (FDA CBER Webinar, 2024).

In this study, we evaluated next-generation sequencing (NGS) strategies for characterizing chemically synthesized gRNAs. We found that the widely used SMARTer assay, while capable of producing sequenceable libraries, introduced substantial artifacts during library preparation. These included truncated scaffold species at oligo(A) stretches in the scaffold region and 5′(n-1) deletions within the spacer sequence. Although absent in the original gRNA, these artifacts accounted for over 10% of the sequencing reads, creating the false appearance of impurities. Through experimental and computational approaches, we traced these artifacts to mispriming by template-switching oligonucleotides (TSOs). Importantly, these artifacts occur during sequencing, and although they do not reflect real gRNA impurities, they compromise assay accuracy and can obscure true sequence impurities.

To overcome these limitations, we developed FUSS-seq (Full-length Uncoupled Second-strand Synthesis followed by sequencing), a novel assay that integrates principles from 5′ RACE with a modified TSO bearing a 3′ polymerase-blocking moiety. FUSS-seq markedly reduced artifacts and increased full-length gRNA recovery, providing a more accurate and lower-bias method for gRNA purity assessment. This approach supports improved Chemistry Manufacturing and Controls (CMC) characterization of gRNAs and strengthens the analytical toolkit needed for reliable CRISPR-based therapeutic development.

## Introduction

On December 8^th^, 2023, the Food and Drug Administration (FDA) approved Casgevy, the first CRISPR-based genome editing therapy, developed by Vertex Pharmaceuticals and CRISPR Therapeutics, for sickle cell disease (SCD) (Frangoul H, 2024). **This milestone underscored the clinical potential of genome editing**. At Vor Bio, CRISPR-Cas9-mediated knockout of the CD33 surface protein was applied to hematopoietic cells derived from healthy donors. These edited cells were then infused into conditioned Acute Myeloid Leukemia (AML) patients (NCT04849910). After bone marrow engraftment, tumor recurrence was treated with CD33-targeted immunotherapies. The healthy donor-derived CD33-negative cells were spared by the treatment, demonstrating that CD33 knockout effectively shielded the healthy cells from the therapeutic agent (Lydeard JR, 2023). Together, these early trials illustrate the clinical promise of CRISPR-based medicines.

CRISPR-based medicines require two components: a guide RNA that specifically directs the gene editing machinery to the target genomic location, and the Cas9 enzyme, an endonuclease capable of generating DNA double-stranded breaks that trigger cellular DNA repair mechanisms that can result in gene knockout (Cho SW, 2013; Cong L, 2013; Jinek M, 2013; Mali P, 2013.). As with other clinical drug substances, both critical components demand extensive Chemistry, Manufacturing, and Controls (CMC) characterization prior to use in Good Manufacturing Practices (GMP) manufacturing (FDA-2024-D-1244; FDA-2024-D-4311).

gRNAs consist of two elements (Supplementary Fig. 1A): (1) A spacer sequence that guides Cas9 to the target DNA; this sequence is typically 20-nucleotide long and is designed to be specific to the desired target DNA sequence; (2) A backbone sequence that is typically 80-nucleotides long and is important for Cas9 binding and catalytic activity (Jinek M, 2012). Chemically synthesized gRNAs are produced by solid-phase oligonucleotide synthesis in the 3′ to 5′ direction and then purified through multiple steps of High-Performance Liquid Chromatography (HPLC) and tangential flow filtration (TFF) to enrich the full-length, 100-mer product. The synthesis and purification steps are not 100% effective and fail to fully remove impurities, which can include shorter sequences (e.g., 1 nucleotide deletion or “n-1”) or base substitutions. These molecules are often modified at the 5′ and 3′ ends to increase stability and prevent enzymatic degradation (Supplementary Fig. 1A)(Hendel A, 2015). The fidelity of the spacer sequence is of critical importance since this element of the gRNA dictates the specificity of the editing machinery, and errors could lead to unintended off-target genomic loci, posing a safety risk to patients. Regulatory guidance recommends ≥80% purity by liquid chromatography (LC) and full characterization of any peaks representing ≥1% of the product (FDA CBER Webinar, 2024). In addition, orthogonal base-level sequence assessment using a sensitive, high-throughput sequencing method, such as next-generation sequencing (NGS), is advised (FDA-2024-D-4311).

Despite this guidance, current sequencing approaches for gRNA characterization remain underdeveloped. Most commercial NGS kits were designed for endogenous small RNAs and do not account for the unique structural features or chemical modifications of synthetic gRNAs. As a result, they can generate technical artifacts that mimic true impurities, leading to inaccurate purity assessments and the potential for misinterpretation in regulatory submissions. Accurate, artifact-free sequencing is therefore a critical unmet need for advancing gRNA quality control and ensuring the safe clinical application of CRISPR therapeutics.

To address this need, we evaluated commercially available NGS short-read sequencing, library preparation kits designed for small RNAs. The small RNA Switching Mechanism at 5′ End of RNA Template (SMARTer) kit from Takara has been previously applied to gRNAs to characterize their sequence identity (Macias LA, 2023; Katta V, 2024). However, we observed that the template-switching mechanism introduces artifacts that compromise accurate purity characterization. To overcome these limitations, we developed a new method, FUSS-seq, which enables more accurate base-level assessment of gRNA purity characterization.

## Results

### Assessing methods for full-length gRNA sequencing

We first evaluated two strategies to generate Illumina-compatible DNA libraries from chemically synthesized gRNAs: (1) ligation-based and (2) template-switching-based methods. The ligation-based approach involved the following key steps (Supplementary Fig. 1B): (i) 5′ phosphorylation of the gRNA; (ii) ligation of a 3′ adapter; (iii) ligation of a 5′ adapter; (iv) reverse transcription using a primer specific to the 3′ adapter; (v) indexing PCR; and (vi) Illumina sequencing. None of the tested commercial kits (see Materials and Methods) yielded a library of the expected size; capillary electrophoresis revealed only primer dimers (Supplementary Fig. 1C). This outcome is consistent with previous reports that chemical modifications on gRNAs can inhibit ligation (Munafó DB, 2010) and suggests that additional steps are required to obtain sequenceable libraries from synthetic gRNAs.

We next tested a template-switching-based method using the smRNA SMARTer kit (Takara). The protocol includes: (i) RNA polyadenylation; (ii) reverse transcription using an oligo(dT) primer with template switching; (iii) indexing PCR; and (iv) Illumina sequencing (Fig. 1A). Initial experiments with a full-length 100-mer gRNA generated by two vendors produced bands of the expected size (Supplementary Fig. 2A and Supplementary Fig. 4A and B). Sequencing confirmed full-length gRNA identity, as shown in other studies (Macias LA, 2023; Katta V, 2024). Analysis of the sequencing data revealed two major impurity species: truncated scaffold (~20%) and 5′(n-1) (typically >10%) variants (Fig. 1B and C). Interestingly, LC of the full-length gRNA did not detect large peaks that would represent the truncated scaffold or the 5′(n-1) species (Supplementary Fig. 3A and Supplementary Fig. 4A and B). While LC resolution of the 5′(n-1) is challenging, >10% is detectable, suggesting that the observed scaffold variants may be NGS artifacts. Sequence analysis showed that the truncated scaffold variants overlapped with A-rich regions about 20 bp to the 3′ end of the gRNA scaffold, indicating unintended binding of the oligo(dT) primer. To mitigate this, we replaced the standard SMARTer oligo(dT) primer with an anchored variant ending in (A)_3_, complementary to the (U)_3_ sequence at the 3′ end of the gRNA. This modification eliminated the truncation artifacts in the scaffold artifacts (Fig. 1B and C).

**Figure 1.**
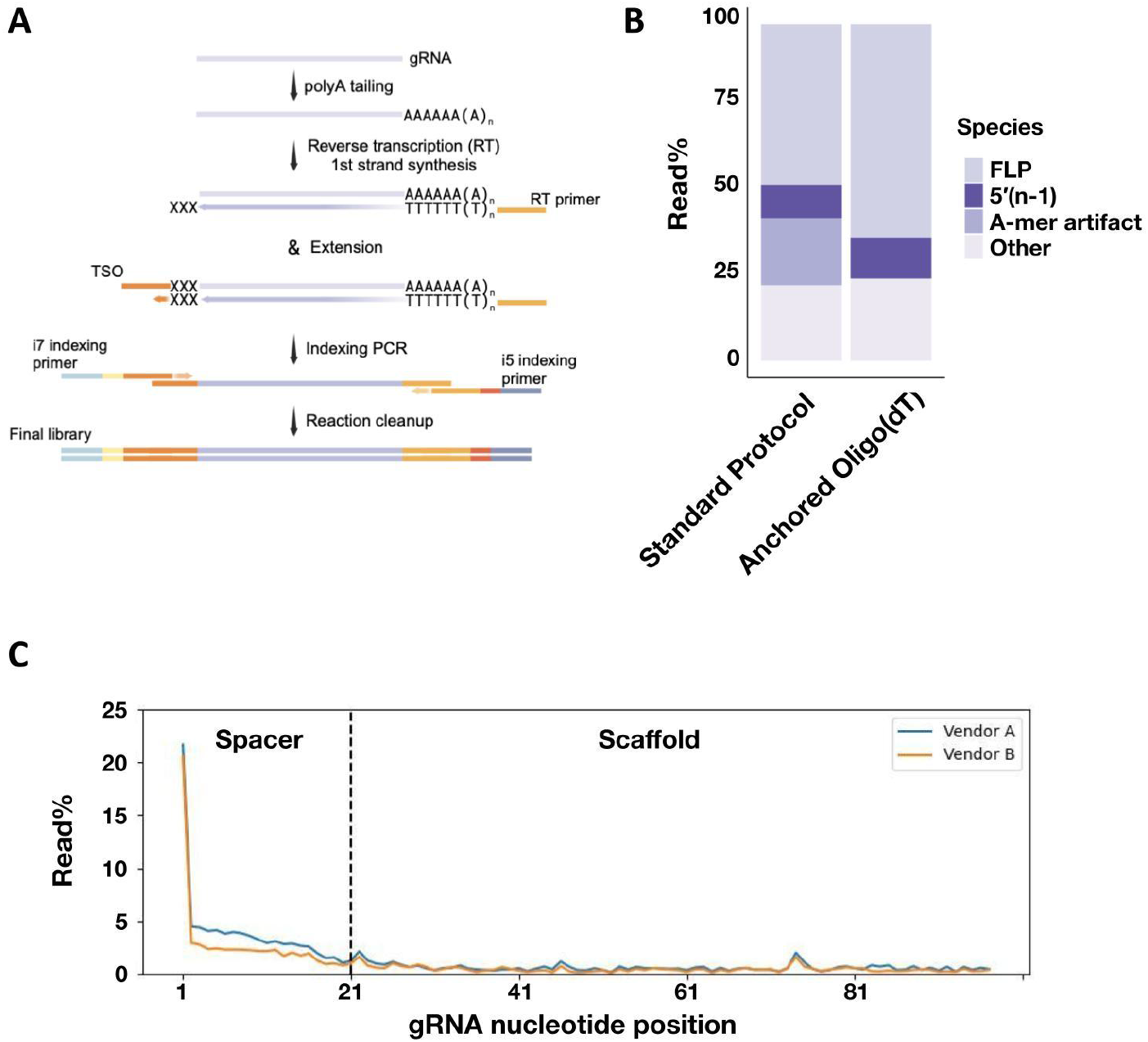
Limitations of the SMARTer protocol introduce artifacts in sgRNA sequencing. **(A)** Schematic overview of the SMARTer protocol highlighting the sequence of reactions and molecular intermediates. **(B)** Stacked barplot comparing the percentages of major species generated by the standard SMARTer protocol or the protocol incorporating a modified anchored oligo(dT) primer. FLP, Full-Length Product. **(C)** Line plot showing the percentage of reads with deletions or mismatches by nucleotide position generated by the SMARTer protocol.

### LC–MS/MS analysis identifies gRNA 5′(n-1) impurities as an NGS artifact

To determine whether the 5′(n-1) impurity was a true synthesis impurity or an artifact of library preparation, we analyzed the gRNA using an independent LC-MS/MS-based approach. To improve analytical resolution, we developed an LC–MS/MS–based RNase H cleavage assay targeting the 5′ region of sgRNAs (Fig. 2A). The assay proceeds as follows: (i) hybridization of the gRNA to an antisense oligonucleotide (ASO) complementary to the conserved repeat near the 5′ end of the scaffold; (ii) RNase H cleavage of the RNA–DNA heteroduplex, generating a short 5′ fragment and a longer conserved fragment; (iii) LC separation of the cleavage products; and (iv) MS-based peak identification. Applying this assay to full-length gRNAs and designed impurity controls revealed that the 5′(n-1) species observed by NGS was indeed a library preparation artifact (Fig. 2B–D). Comparing the LC peaks of the 10% spike-in 5′(n-1) impurity control with the full-length gRNA demonstrated that this species is present at a lower rate in the full-length gRNA than the >10% reported when using the SMARTer kit (Fig. 2C). LC followed by mass spectrometry analyses showed that the 5′(n-1) impurity was less than 0.5% of the total peaks (Fig. 2D). These results confirm that the 5′(n-1) signal in sequencing data primarily arises from technical artifacts rather than true gRNA impurities.

**Figure 2.**
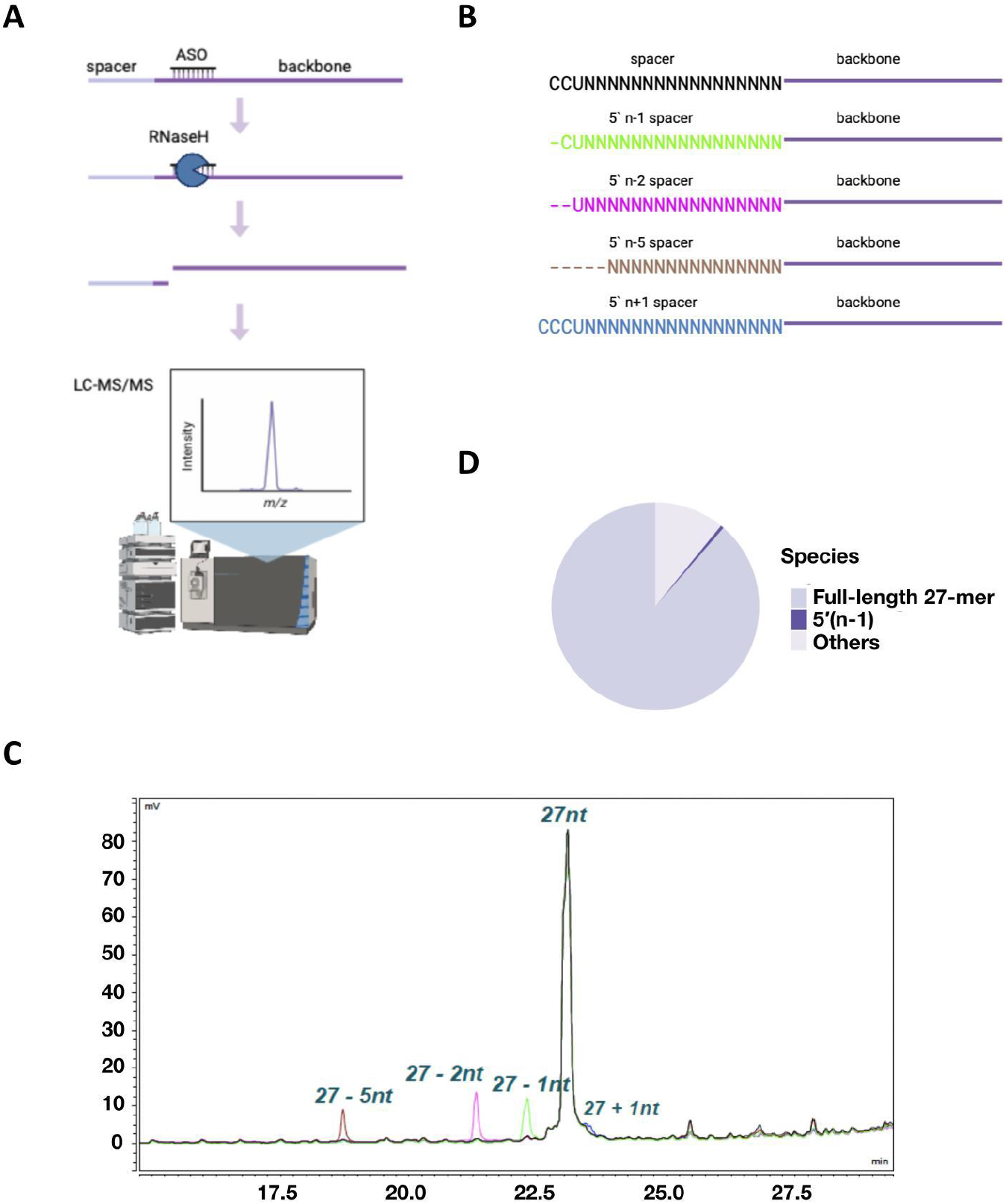
The 5′(n-1) peak is a technical artifact from SMARTer, not a synthesis impurity. **(A)** Schematic representation of the LC-MS/MS strategy used for gRNA impurities characterization as an orthogonal assay to NGS. **(B)** Synthetic spike-in gRNAs used to trace gRNA impurities. **(C)** LC Chromatogram traces (UV) showing 27-mer gRNA products generated by the RNase H, and associated spike-ins. **(D)** Pie chart displaying area percentages of LC-MS/MS identified species.

### Template-switching bias drives sequence-dependent 5′(n-1) artifacts

To explore the source of this artifact, we first designed synthetic PCR and sequencing controls that mimic the last two steps of the SMARTer protocol (Supplementary Table S1). Sequencing of these controls revealed the 5′(n-1) species at a < 0.1% read frequency, localizing the artifact to the reverse transcription step (Fig. 3A). We then asked if the artifact was specific to template-switching-based protocols or was a general reverse-transcriptase issue. We analyzed published datasets comparing template-switching and ligation-based methods (Dard-Dascot C., 2018). In human miRNA samples, the SMARTer kit exhibited a higher frequency of 5′(n-1) artifacts than the TruSeq ligation method (Fig. 3B, Supplementary Fig. 5A). These results indicate that the artifact is specific to the template-switching mechanism rather than reverse transcription *per se*.

**Figure 3.**
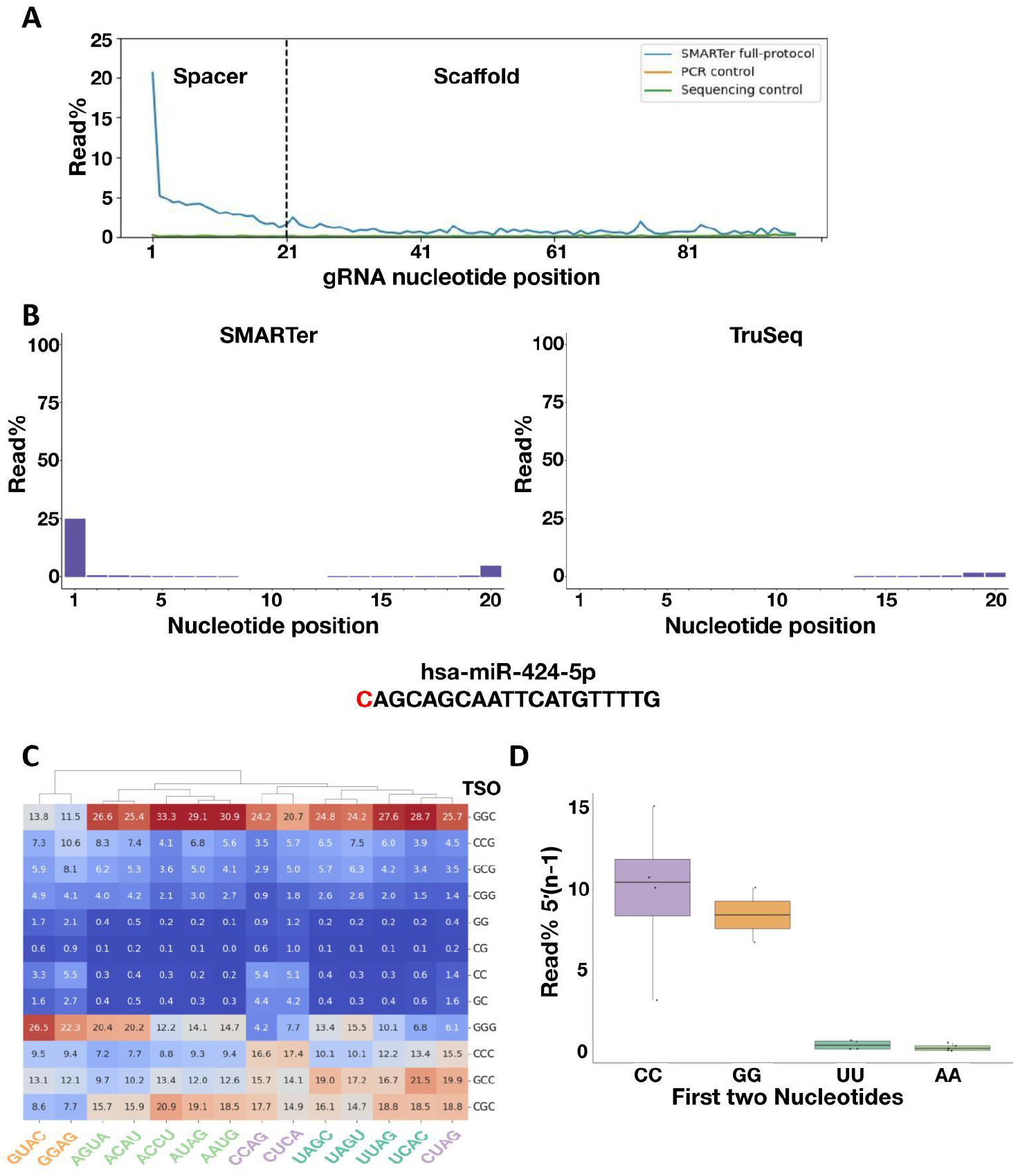
The 5′(n-1) artifact originates during template-switching oligo. **(A)** Line plot showing the percentage of reads with deletions or mismatches at each nucleotide position, comparing the SMARTer product to PCR and sequencing controls. **(B)** Bar plot showing the percentage of reads with deletions at each nucleotide position of miRNA hsa-miR-424-5p (first 20 nucleotides), comparing SMARTer to TruSeq. **(C)** Heatmap displaying the incorporation of SMARTer TSO sequences across gRNAs of different 5’-end nucleotides. **(D)** Box and whisker plot showing the percentage of the 5′(n-1) truncation species in relation to the 5′ nucleotides in the gRNA sequence. The box represents the interquartile range; the line represents the median; whiskers represent the upper and lower limits within 1.5x the interquartile range.

Moloney murine leukaemia virus (MMLV)-derived reverse transcriptases add non-templated nucleotides, often cytosines, at the 3′ end of cDNA upon reaching the 5′ end of the RNA (Clark JM, 1988; Schmidt WM, 1999). These non-templated additions enable the TSO 3′ end to anneal and facilitate template switching (Zhu YY, 2001). Analysis of TSO sequences revealed that the majority of additions were GC-rich 3-nucleotides, accounting for 82-99% of reads depending on the gRNA 5′ leading nucleotide identity (Fig. 3C). Strikingly, gRNAs starting with cytosines were associated with high percentages of 2-nucleotide additions (~10% median, Fig. 3D). This coincides with the SMARTer-derived 5′(n-1) artifacts observed, suggesting a sequence-dependent mispriming bias. To test this, we synthesized semi-identical gRNAs that differed only in the first two 5′ nucleotides. These synthetic gRNAs started with CC, GG, UU, or AA. The CC and GG variants showed elevated n-1 levels (Fig. 3D), supporting the hypothesis that TSO mispriming on 5′-terminal Cs or Gs drives artifact formation.

### FUSS-seq: A 5′-End Capture Strategy for Accurate gRNA Spacer Profiling

To mitigate the artifacts generated by TSO mispriming events, we hypothesized that developing an assay designed to capture the 5′ end of RNA molecules would greatly improve the purity characterization of the gRNA spacer region. Inspired by the 5′ Rapid Amplification of cDNA Ends (RACE) workflow (Frohman MA, 1988), we developed a new assay called Full-length Uncoupled Second-strand Synthesis followed by sequencing (FUSS-seq). Our method proceeds as follows (Fig. 4A): (i) polyadenylation of gRNA; (ii) reverse transcription with a reverse transcriptase lacking RNase activity; (iii) template switching to the TSO; (iv) RNase H degradation of RNA, including the rNrNrN tail of the TSO; (v) second-strand synthesis primed by the DNA portion of the TSO; and (vi) PCR amplification for indexing and sequencing.

**Figure 4.**
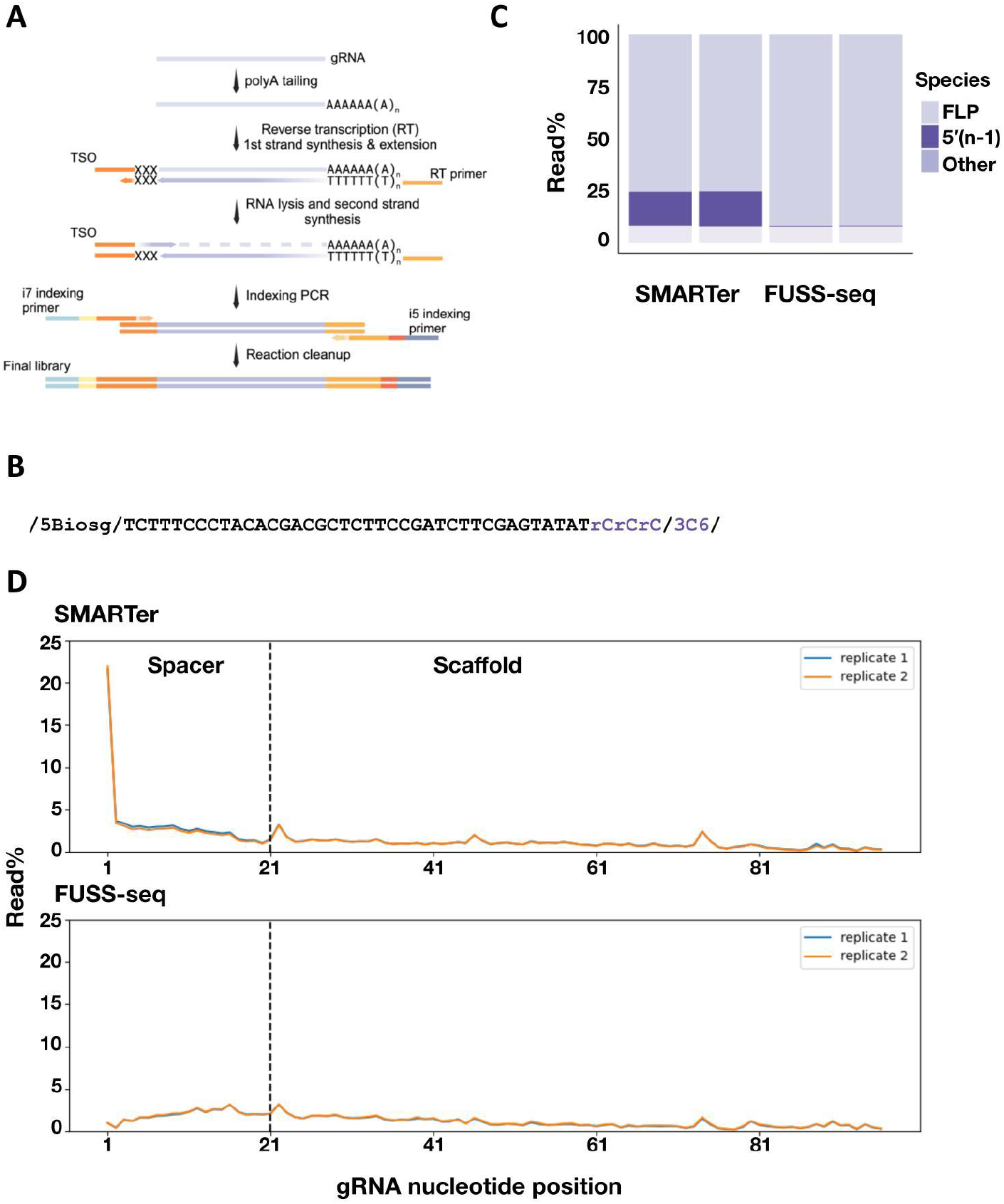
FUSS-seq eliminates the 5′(n-1) issue by redesigning the TSO. **(A)** Schematic representation of the FUSS-seq protocol, with a specially engineered 3′-end TSO to reduce template-switching errors. **(B)** Detailed design of the FUSS-seq TSO. The major TSO FUSS-seq modifications, the ribo-cytosine tracts (rCrCrC) and 3′ hexanediol moiety (3C6), are highlighted in violet. **(C)** Stacked barplot comparing SMARTer and FUSS-seq. The two bars per protocol indicate two technical replicates. FLP, Full-Length Product. **(D)** Line plot showing level of deletions or mismatches by nucleotide position in SMARTer (top) and FUSS-seq (bottom).

A critical component of FUSS-seq is the modified TSO, which contains a fixed rCrCrC 3′ sequence. By contrast, the SMARTer TSO contains variable terminal bases (Fig. 3C), and most published TSOs use rGrGrG (Zhu YY, 2001). While rGrGrG reduces 5′(n-1) artifacts, it introduces other unexpected species, particularly in the 5′ (2-15bp) region (Supplementary Fig. 6A and B). This first version of the FUSS-seq rCrCrC TSO produced a moderate level of 5′(n-1) artifacts (~4%), less than SMARTer (~10–15%) but higher than rGrGrG (Supplementary Fig. 6A and B), indicating partial mitigation via the 5′ RACE workflow.

In the second-strand synthesis step of the 5′ RACE workflow, RNA in RNA-cDNA duplexes is degraded by RNaseH. Degradation also affects the 3′ ribonucleotides of the TSO oligo. A second strand of DNA is then synthesized, after a short denaturation step, by a DNA polymerase, where the TSO functions as the primer. We speculated that some of this residual mispriming might happen during the second-strand synthesis step. After RNaseH degradation, we will have two TSO species: (1) Ribo-depleted TSO molecules, which are missing the 3′ ribonucleotides; Full-length TSO molecules, which did not participate in the reverse transcription, did not hybridize to cDNA, and, therefore, did not undergo RNA degradation of the 3′ ribonucleotides. Full-length TSO might be a considerable percentage of molecules since it is provided in high quantities during reverse transcription. During DNA polymerization, these longer TSOs might erroneously misprime to the cDNA and lead to artifacts like the observed 5′(n-1) species. We hypothesized that residual full-length TSOs, present in excess and not degraded by RNaseH, could misprime during second-strand synthesis. To prevent these events, we generated TSO oligos carrying a hexanediol blocking moiety in the last 3′ nucleotides (Fig. 4B). This nucleotide modification does not prevent mispriming events but hinders polymerization if these unwanted events occur. Library preparation with this modified TSO oligo resulted in a great reduction of the 5′(n-1) species (Fig. 4C). Moreover, we observed an increase in the percentage of full-length gRNA sequences similar to the LC-MS/MS results (~90%), suggesting that our method improves the overall quality of library preparation (Fig. 4D).

## Discussion and Conclusion

Accurate gRNA purity characterization is a critical component of CMC for CRISPR-based therapeutics. While LC offers a robust method for bulk analysis, it lacks the sensitivity to detect low-abundance variants or confirm sequence identity. NGS offers the throughput and base-level resolution necessary to complement LC-based methods, but, as our results reveal, its accuracy is heavily influenced by library preparation artifacts.

Our evaluation of commercially available small RNA NGS kits highlighted major limitations when applied to chemically synthesized gRNAs. The template-switching-based SMARTer method, although capable of producing full-length sequencing libraries, introduced prominent artifacts, most notably 5′(n-1) variants and truncations in the scaffold region. Through experimental and computational analyses, we traced the origin of these species to TSO mispriming events during reverse transcription and second-strand synthesis.

These findings underscore the importance of method-specific biases in RNA sequencing workflows, particularly for structured, chemically modified, or synthetic RNAs like gRNAs. Misinterpretation of TSO-induced artifacts as genuine synthesis impurities could compromise regulatory submissions or lead to inaccurate assessments.

To overcome these limitations, we developed FUSS-seq, a novel approach that combines key aspects of the 5′ RACE methodology with a modified TSO bearing a 3′ polymerase-blocking moiety. This strategy reduces the likelihood of polymerase extension from residual, misprimed TSOs and significantly improves the fidelity of 5′ gRNA capture. Compared to the SMARTer protocol, FUSS-seq showed markedly reduced artifact levels and improved recovery of full-length gRNA sequences, supporting its utility as a more accurate and lower-bias method for gRNA purity assessment that could be incorporated into a panel of analytical assays suitable for batch release.

Importantly, FUSS-seq is modular and adaptable. Alternative reverse transcriptases, RNA-stabilizing modifications, or orthogonal ligation-free capture strategies could be integrated to expand the method’s applicability to different RNA formats or chemistries. Beyond Cas9-based gRNAs, FUSS-seq might be deployed on other CRISPR modalities, including Cas12 and Cas13 enzymes, as well as prime editing (Zetsche B, 2015; Abudayyeh OO, 2016; Anzalone AV, 2020).

In conclusion, FUSS-seq enhances the analytical toolkit available for the sequence-level characterization of gRNAs, a key requirement for the regulatory success of CRISPR-based gene editing therapeutics. It fills a critical gap in current CMC workflows by minimizing sequencing artifacts and improving full-length product detection. Its implementation can streamline regulatory compliance, improve batch-to-batch consistency, and ultimately support safer and more reliable genome-editing therapies.

## Materials and Methods

### gRNA library preparation with a ligation-based kit

Library preparations using the NEBNext Small RNA Library Prep Set for Illumina (NEB, #E7330) protocol were carried out starting from 500 ng gRNA and according to the manufacturer’s instructions. The following modifications to the protocol were applied: (1) 3,000 ng of synthetic gRNA were phosphorylated using the T4 Polynucleotide Kinase (NEB, #M0201) by incubating the reaction at 37°C for 60 minutes and the enzyme heat-inactivated at 65°C for 20 minutes; (2) 5 ul of reactions were used as input to the library prep; (3) A single, double-sided size selection (0.65x, 1x) with SPRIselect DNA Size Selection Reagent (Beckman, #B23317) was performed to streamline the protocol. Library prep success was tested by running capillary electrophoresis with the TapeStation 4200 DNA High-Sensitivity D5000 kit (Agilent, #5067-5593).

### gRNA library preparation with SMARTer

Library preparations using the SMARTer smRNA-Seq Kit for Illumina (Takara Bio, #635029) protocol were carried out starting from 500 ng gRNA and according to the manufacturer’s instructions. The following modifications to the protocol were applied: (1) An anchored oligo(dT) primer (Supplementary Table S1) was designed to reduce the observed polyA artifacts (see the Results section); (2) A total of 15x PCR cycles were used. (3) A single, double-sided size selection (0.65x, 1x) with SPRIselect DNA Size Selection Reagent (Beckman, #B23317) was performed to streamline the protocol. Library prep success was tested by running capillary electrophoresis with the TapeStation 4200 DNA High-Sensitivity D5000 kit (Agilent, #5067-5593).

### gRNA purity assessment by HPLC

Mobile phases were prepared as follows:

Mobile Phase A:

1. Add the reagents in order to a glass bottle:
  a. 453 ml LC-MS grade water (Fisher Scientific, #6003078),
  b. 5% Methanol HPLC grade (Sigma Aldrich, #34860),
  c. 400 mM Hexafluoroisopropanol (TCI, #0424),
  d. 18 mM Triethylamine (Acros, #21951-0500).
2. Close bottle tightly with cap and seal with parafilm.
3. Shake vigorously for ~ 1 minute.
4. Visually inspect for any undissolved reagents and shake for another 1 minute if any undissolved material remains.

Mobile Phase B:

1. Add the reagents in order to a glass bottle:
  a. 178 ml LC-MS grade water (Fisher Scientific, #6003078),
  b. 60% Methanol HPLC grade (Sigma Aldrich, #34860),
  c. 400 mM Hexafluoroisopropanol (TCI, #0424),
  d. 18 mM Triethylamine (Acros, #21951-0500).
2. Close bottle tightly with cap and seal with parafilm.
3. Shake vigorously for ~ 1 minute.
4. Visually inspect for any undissolved reagents and shake for another 1 minute if any undissolved material remains.

Synthetic gRNA was diluted to 0.5 mg/ml in nuclease-free water (Invitrogen, #AM3397) and run on a 1290 Infinity III LC System (Agilent) with the following conditions:

1. Flow rate, 0.5 mg/ml,
2. Injection volume, 10 ul,
3. Column temperature, 80°C,
4. Detection, 260 nm,
5. Acquisition rate, 10 Hz.

| Time<br>(Min) | MP A<br>(%) | MP B<br>(%) |
| --- | --- | --- |
| 0.0 | 77 | 23 |
| 2.0 | 76 | 24 |
| 24.0 | 69 | 31 |
| 28.0 | 69 | 31 |
| 30.0 | 64 | 36 |
| 31.0 | 0 | 100 |
| 38 | 0 | 100 |
| 39 | 77 | 23 |
| 45 | 77 | 23 |

### gRNA 5′-end characterization with an LC/MS-based RNaseH cleavage assay

The 5′-end of the gRNA was fragmented and analyzed using an LC-MS/MS-based RNaseH cleavage assay developed by AxoLabs. An antisense oligo (ASO) targeting the conserved crRNA repeat region downstream of the spacer was designed and optimized for RNaseH cleavage of the 5′-end of the gRNA. The cleavage fragments are subjected to IP-RP-HPLC for separation, detection, and quantification using a UV detector. The HPLC is followed by ESI-MS characterization using an ESI-qTOF mass spectrometer. After a blank subtraction, HPLC peaks are reported with retention time and relative peak area. The molecular weights observed from the ESI-MS are reported for each peak, from which impurity characterization is conducted. The ASO design, RNaseH cleavage, and HPLC conditions were optimized for the gRNA sequence. For quantifying gRNA truncated impurities by HPLC, the UV detector was calibrated with a dilution series of full-length gRNA, resulting in a relative quantification of >1% peak area.

### gRNA library preparation with FUSS-seq

Perform the FUSS-seq protocol as follows:

1. gRNA dilution: Dilute gRNA to 2,500 ng in 6.5 ul of molecular-grade H2O.
2. Polyadenylation:
  a. Prepare the mix on ice as follows (volumes for one reaction):
    i. 2 ul E. coli Poly(A) Polymerase Reaction Buffer (10x)(NEB, #M0276),
    ii. 1 ul ATP (10 nM)(NEB, #M0276),
    iii. 0.25 ul E. coli Poly(A) Polymerase (5 U/ul)(NEB, #M0276),
    iv. 0.25 ul molecular-grade H2O,
  b. Add 3.5 ul of reaction to 6.5 ul of gRNA (2,500 ng), mix on ice, and immediately incubate at 37°C for 30 minutes.
  c. Transfer the sample on ice and proceed immediately with the next step.
3. Oligo(dT) annealing:
  a. Prepare the mix on ice as follows (volumes for one reaction):
    i. 2 ul molecular-grade H2O,
    ii. 1 ul dNTPS (10 mM) (NEB, #N0447),
    iii. 1 ul anchored oligo(dT) primer (Supplementary Table S1),
  b. Add 4 ul of reaction to 2 ul of polyadenylated gRNA, mix on ice, immediately incubate at 70°C for 5 minutes.
  c. Transfer the sample on ice and proceed immediately with the next step.
4. Reverse transcription:
  a. Note: Preparation of the reverse transcription mix during the annealing step is recommended.
  b. Prepare the mix on ice as follows (volumes for one reaction):
    i. 2.5 ul Template Switching RT Buffer (4x) (NEB, #M0466),
    ii. 0.5 ul FUSS-seq TSO (75 uM) (Supplementary Table S1),
    iii. 1 ul Template Switching RT Enzyme Mix (10x) (NEB, #M0466).
  c. Add 4 ul of reaction to the annealed gRNA, mix on ice, and immediately incubate at 42°C for 90 minutes. Heat-inactivate at 85°C for 5 minutes.
  d. Transfer the sample on ice and proceed with the next step.
5. Second-strand synthesis:
  a. Prepare the mix on ice as follows (volumes for one reaction):
    i. 62 ul molecular-grade H2O,
    ii. 20 ul Q5U buffer (5x) (NEB, #M0515),
    iii. 2 ul dNTPs (10 mM) (NEB, #N0447),
    iv. 1 ul Q5U DNA polymerase (2 U/ul) (NEB, #M0515),
    v. 5 ul E. coli RNase H (5 U/ul) (NEB, #M0297),
  b. Add 90 ul of mix to 10 ul of reverse-transcribed gRNA and mix on ice.
  c. Incubate at 37°C for 30 minutes, 95°C for 1 minute, 55°C for 1 minute, and 65°C for 10 minutes.
  d. Proceed with double-sided size selection (0.65x, 1.00x) with the SPRIselect DNA Size Selection Reagent (Beckman, #B23317). Elute in 20 ul molecular-grade H2O.
6. Indexing PCR:
  a. Prepare the mix on ice as follows (volumes for one reaction):
    i. 26 ul molecular-grade H2),
    ii. 50 ul Q5 master-mix (2x) (NEB) (M0492),
    iii. 4 ul xGen™ UDI 10nt Primer Plates 1-4 (IDT, #10008052).
  b. Add 80 ul of mix to 20 ul of eluted double-stranded cDNA and mix on ice.
  c. Incubate at 98°C for 5 minutes, 98°C for 10 seconds, 65°C for 10 seconds, 72°C for 20 seconds, repeat these short steps 14x (total 15x cycles), 72°C for 2 minutes.
  d. Proceed with double-sided size selection (0.65x, 0.90x) with the SPRIselect DNA Size Selection Reagent (Beckman, #B23317). Elute in 20 ul molecular-grade H2O.
  e. Library prep success was tested by running capillary electrophoresis with the TapeStation 4200 DNA High-Sensitivity D5000 kit (Agilent, #5067-5593).
  f. Proceed with Qubit quantification (ThermoFisher Scientific, #Q33230) and Illumina sequencing.

### gRNA libraries sequencing

SMARTer and FUSS-seq libraries were sequenced on the Illumina MiSeq instrument (SY-410-1003) with the MiSeq Reagent Kit v2 300-cycles (Illumina, #MS-102-2002). Libraries were loaded at 8 pM with 40% PhiX. The following cycles were used: Read1 121 + Index1 8 + Index2 8 cycles.

### Bioinformatics analysis

SMARTer and TruSeq miRNA data were processed as previously described (Dard-Dascot C, 2018). Briefly, adapters were trimmed, and reads were aligned to the database of small human non-coding RNAs (DASHR, Leung YY, 2016). Aligned reads were then re-aligned against the small RNA reference sequence using a pairwise alignment module (Cock PJA, 2009).

To analyze gRNA sequencing data, poly-A tails were removed using Cutadapt (Martin M, 2011). For SMARTer-derived gRNA sequencing data, the first 3 nucleotides at the 5′-end were removed as TSO following the manufacturer’s recommendation. The resulting sequences were then aligned against the gRNA reference sequence (spacer + scaffold) using a pairwise alignment module (Cock PJA, 2009). We calculated deletions in the 81~97 bp region (starting with the A-mers in the scaffold region). Reads are discarded if there are >17bp deletions in this region (mostly caused by poly-T oligo mispriming). Alignments with excessive mismatches and gaps (>30 bps overall) were further discarded as artifacts.

The FUSS-seq gRNA sequencing data were processed similarly, except that the full-length cleaned NGS reads (including TSO) were aligned against the gRNA reference (TSO + spacer + scaffold). Additionally, alignments with >4bp gaps within the TSO region were also discarded. The remaining alignments were then subjected to TSO trimming, retaining only gRNA alignment (spacer + scaffold). For each NGS dataset, the reads corresponding to each alignment species were counted and used for purity and impurity quantification.

## Data availability

SMARTer and TruSeq miRNA data (Dard-Dascot C, 2018) are available in the Sequence Read Archive (SRA) database, IDs SRR6466802, SRR6466803, SRR6466813, SRR6466814. Data produced at Vor are available upon request.

## Acknowledgments

We thank our colleagues at Vor Bio for their invaluable technical support and insightful contributions towards this study. Figures 1A, 2A, 2B, 4A, and Supplementary Figures 1A and 1B were created using BioRender.com.

## Notes

### Competing Interest Statement

A.M, H.Q., M.S., L.K., S.L., J.H., T.C., H.G.G., J.R.L., R.W., and E.G.A. are
salaried employees of Vor Biopharma and may hold equity in the company.

### Summary of Updates

One of the authors' names was mispelled. This new version has the correct name.

