## Supplementary Material for "Cutting Through the Artifacts: Dissecting gRNA Impurities with FUSS-seq"

#### Supplementary Figure 1

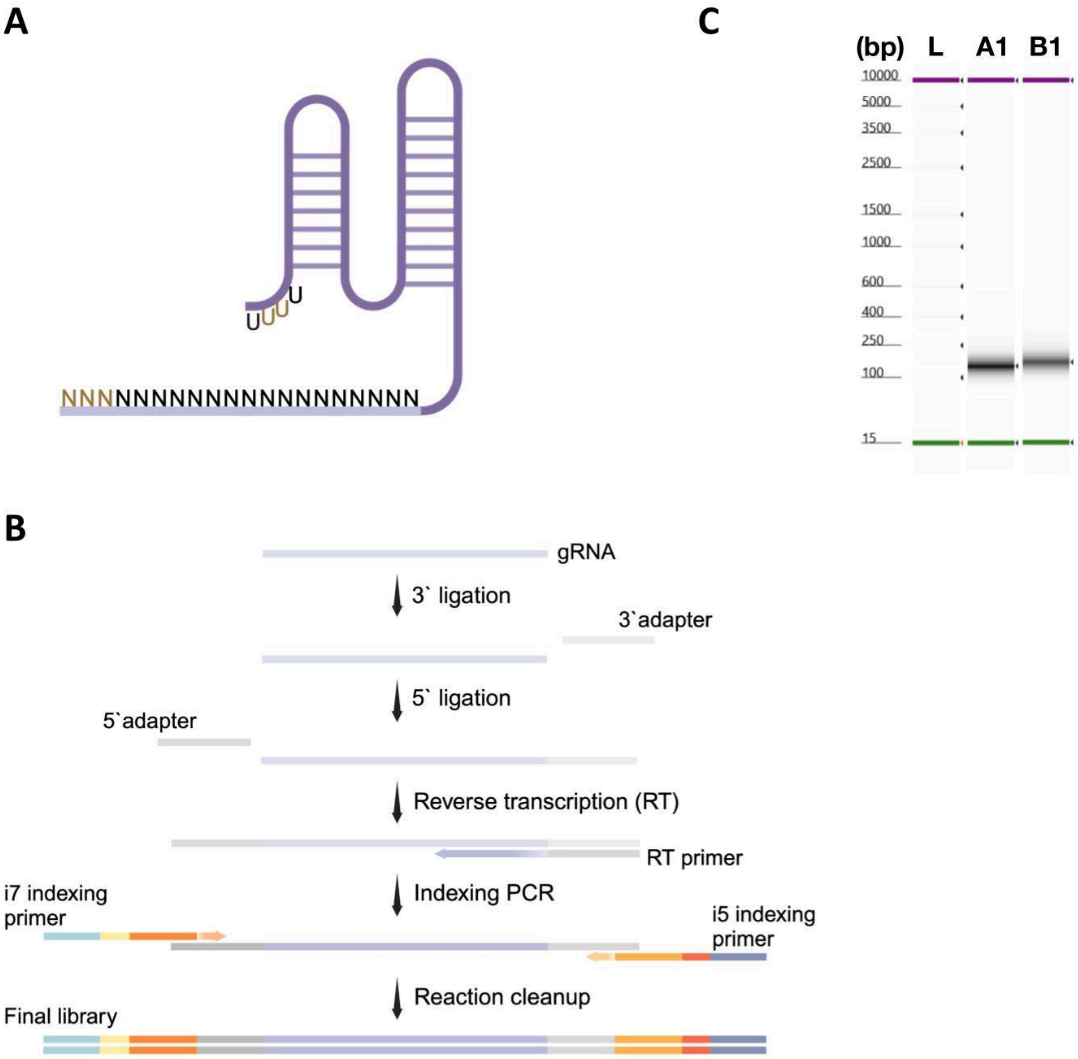

**Supplementary Figure 1. Ligation-based small-RNA library prep did not work in our lab.**
**(A)** Schematic of a gRNA. The letter N indicates any RNA nucleotide (A, U, G, or C). The light-purple line (Ns nucleotides) represents the spacer sequence. The dark-purple line represents the scaffold. Brown nucleotides indicate 2′-O-methyl-ribonucleotides.
**(B)** Schematic overview of the ligation-based protocol, highlighting the sequence of reactions and molecular intermediates.
**(C)** Tapestation electrophoresis showing the size (bp) of libraries generated with a ligation-based kit. Sizes were lower than expected (~250 bp), indicating that the library prep was not successful. L, TapeStation HS D5000 Ladder. A1, gRNA Vendor A. B1, gRNA Vendor B.

### Supplementary Figure 2

A

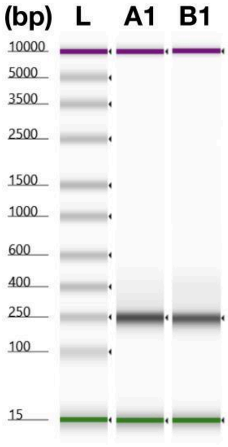

**Supplementary Figure 2. The SMARTer standard protocol generates sequenceable libraries from synthetic gRNAs.**
**(A)** Tapestation electrophoresis showing the size (bp) of libraries generated with the SMARTer kit. Sizes were as expected (~250 bp), indicating that the library prep was successful. L, TapeStation HS D5000 Ladder. A1, gRNA Vendor A. B1, gRNA Vendor B.

#### Supplementary Figure 3

**A**

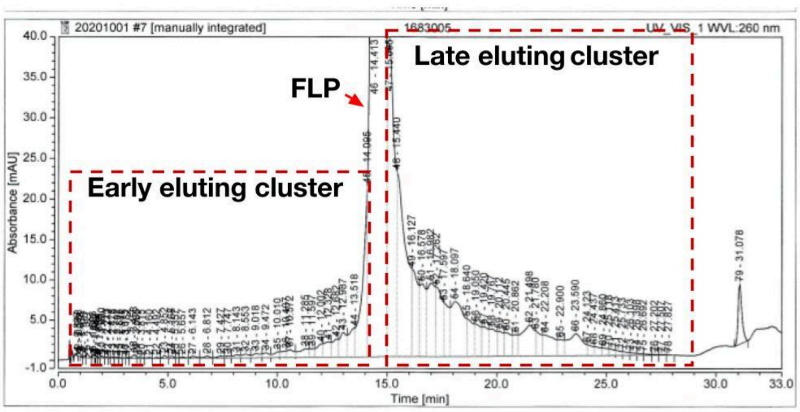

**Supplementary Figure 3. Liquid chromatography analysis of synthetic gRNAs did not reveal any major truncated species.**
**(A)** Liquid chromatography of a synthetic gRNA. FLP, Full-Length Product.

### Supplementary Figure 4

## A

**Vendor A**

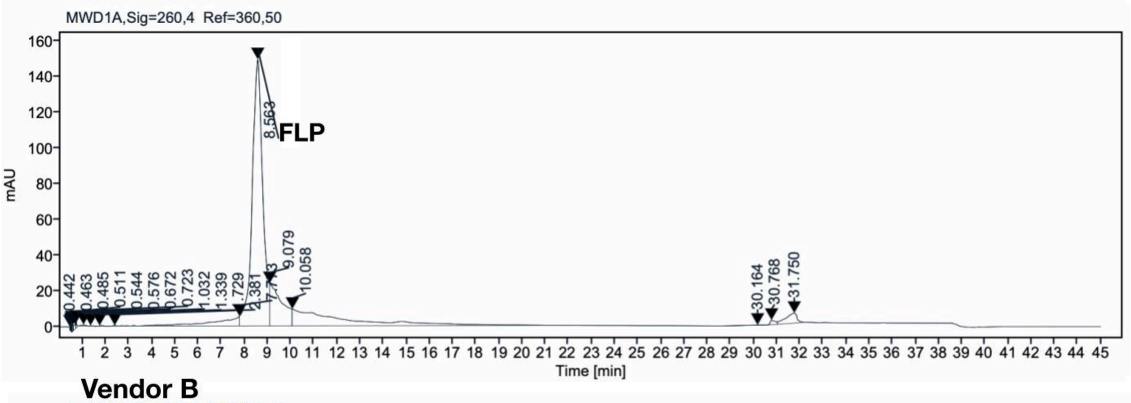


**Vendor B**

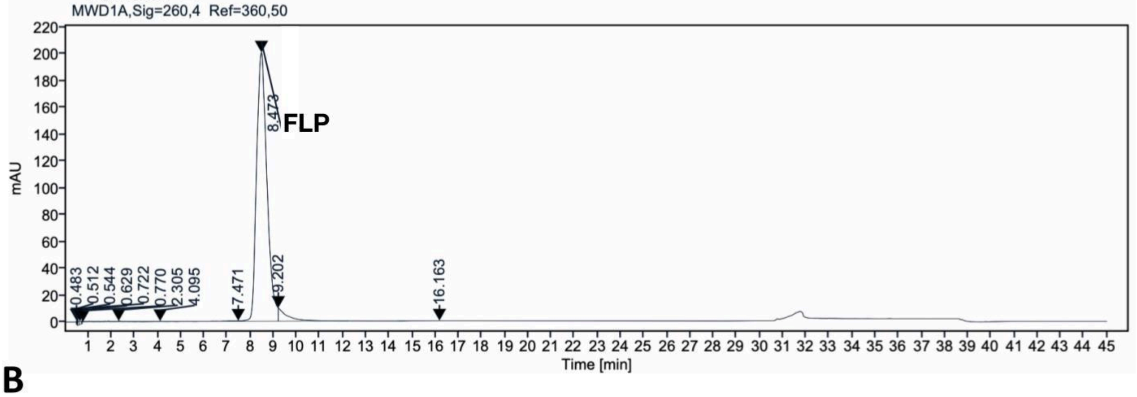


## B

**Vendor A**

| RT (min) | Area (%) |
|---|---|
| 0.442 | 0.04 |
| 0.463 | 0.02 |
| 0.485 | 0.06 |
| 0.511 | 0.08 |
| 0.544 | 0.07 |
| 0.576 | 0.1 |
| 0.672 | 0.19 |
| 0.723 | 0.1 |
| 1.032 | 0.1 |
| 1.339 | 0.03 |
| 1.729 | 0.06 |
| 2.381 | 0.14 |
| 7.773 | 4.92 |
| 8.563 | 60.92 |
| 9.079 | 11.31 |
| 10.058 | 18.77 |
| 30.164 | 0.06 |
| 30.768 | 0.51 |
| 31.75 | 2.5 |

**Vendor B**

| RT (min) | Area (%) |
|---|---|
| 0.483 | 0.05 |
| 0.512 | 0.08 |
| 0.544 | 0.2 |
| 0.629 | 0.18 |
| 0.722 | 0.08 |
| 0.77 | 0 |
| 2.305 | 0.47 |
| 4.095 | 0.27 |
| 7.471 | 0.3 |
| 8.473 | 94.2 |
| 9.202 | 4.04 |
| 16.163 | 0.14 |

**Supplementary Figure 4. Liquid chromatography analysis of synthetic gRNAs did not reveal any major truncated species.**

**(A)** Liquid chromatography profile of Vendor A (top) and B (bottom) synthetic gRNAs. FLP, Full-Length Product.

(**B**) Tables reporting peak area percentages from Vendor A (left) and Vendor B (right). RT, Retention Time.

#### Supplementary Figure 5

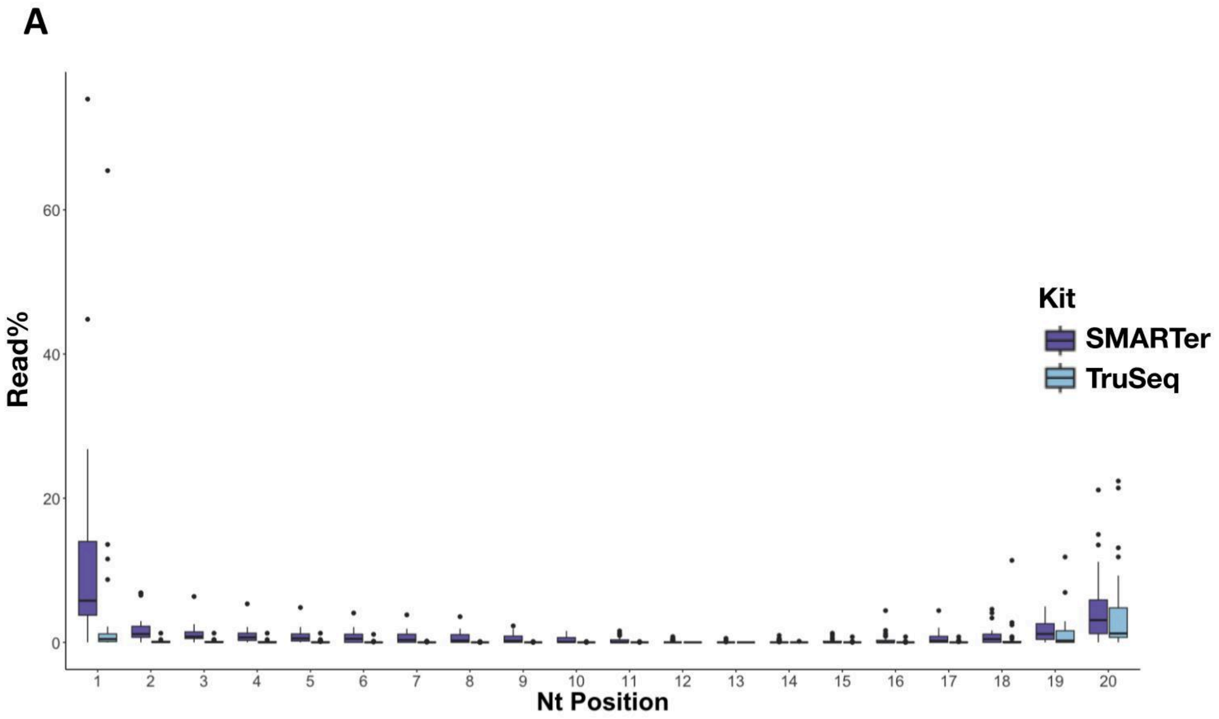

**Supplementary Figure 5. The SMARTer kit shows higher percentages of 5′(n-1) when compared to a ligation-based kit.**

**(A)** Box and whisker plot comparing the percentage of reads with deletions at each nucleotide position (first 20 nucleotides) generated by the SMARTer or the TruSeq (ligation-based kit) protocol. The box represents the interquartile range; the line represents the median; whiskers represent the upper and lower limits within 1.5x the interquartile range.

#### Supplementary Figure 6

**A**

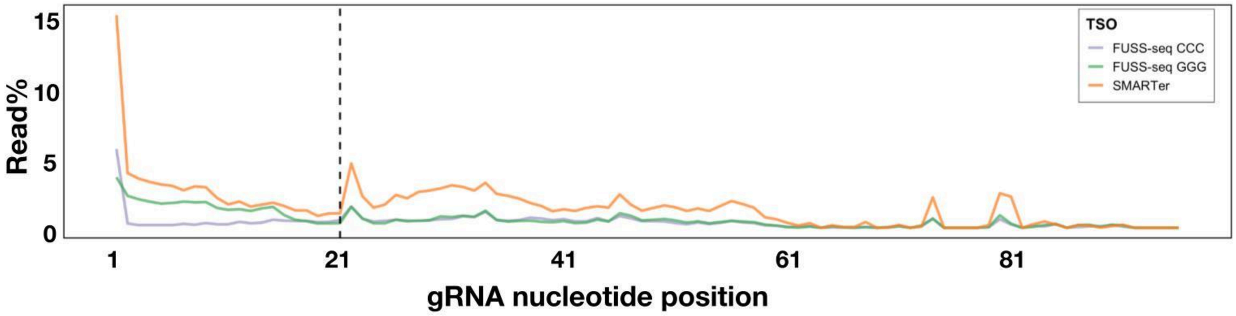


**B**

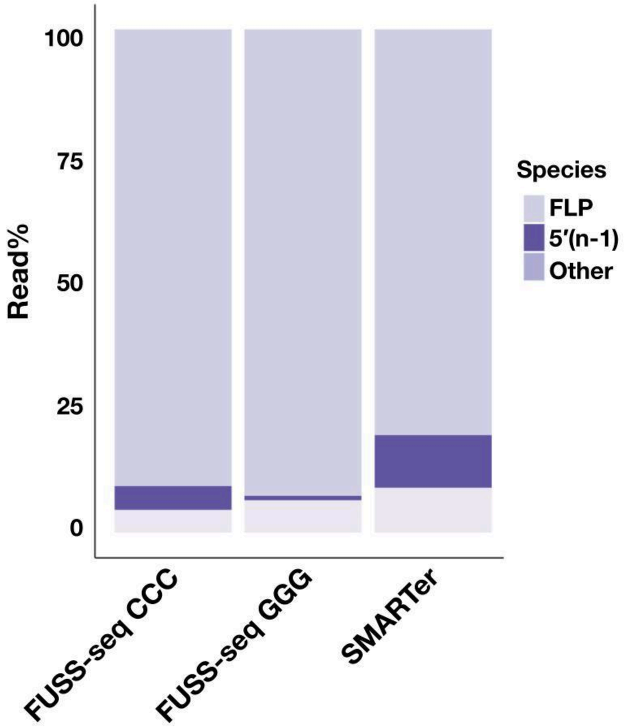

**Supplementary Figure 6. FUSS-seq TSO generates fewer artifacts than the classical TSO ending in rGrGrG.**

**(A)** Line plot showing the percentage of reads with deletions by nucleotide position generated by the SMARTer and the FUSS-seq protocols with the rGrGrG or rCrCrC TSO. **(B)** Stacked barplot comparing SMARTer and FUSS-seq protocols with the rGrGrG or rCrCrC TSO, showing the depletion of n–1 peaks in the FUSS-seq output.
